# ORP2 regulates free cholesterol accumulation in hepatocytes during MASH

**DOI:** 10.1101/2025.01.03.631227

**Authors:** Jin Wu, Yudi Zhao, Liwen Qiu, Yan Liang, Qiaoli Chen, Xiaowei Wang, Jingwen Gu, Hong-Yu Wang, Yang Liu, Xiaoqin Wu, Shuai Chen, Feng-Jung Chen, Mingming Gao, Hongyuan Yang

## Abstract

**Background:** Cholesterol crystals in hepatocytes are known to strongly associate with human Metabolic Dysfunction-Associated Steatohepatitis (MASH). However, it remains unclear which molecular pathway(s) regulates free cholesterol accumulation, and therefore the formation of cholesterol crystals in hepatocytes. In cultured cell lines, oxysterol-binding protein-related protein 2 (ORP2) functions to deliver cholesterol to the plasma membrane from internal organelles.

**Methods:** Here, we generated liver-specific ORP2 knockout (ORP2-LKO) mice and characterized their metabolic phenotypes on chow and high fat diet.

**Results:** The ORP2-LKO mice developed much more severe hepatic steatosis than wild type mice after high-fat diet feeding. They also demonstrated more severe liver inflammation and damage. Notably, free but not esterified cholesterol, as well as cholesterol crystals accumulated in ORP2-LKO liver. The expression of Cyp7a1 was significantly upregulated in ORP2-LKO liver, accompanied with accumulation of taurocholic acid. Our results thus unveil an important *in vivo* function of ORP2 in preventing free cholesterol from accumulating in mouse liver.

**Conclusions:** our results suggest that impaired cholesterol trafficking may enhance the deposition of cholesterol crystals in hepatocytes, promoting the development of MASH.

## Introduction

Cholesterol is an essential constituent of organellar membranes of mammalian cells [1–4]. Cholesterol is synthesized in the endoplasmic reticulum (ER), however, the concentration of cholesterol in the ER is very low (less than 5% of total ER lipids). Instead, up to 90% of cellular free cholesterol exists in the plasma membrane (PM) where it plays a critical role in maintaining the stability and function of the PM [5]. Endosomal compartments also have a relatively high concentration of cholesterol. Thus, the distribution and therefore trafficking of cellular cholesterol are tightly controlled. The oxysterol binding protein (OSBP) and OSBP-related proteins (ORP, for OSBP-related protein) have emerged as key mediators/regulators of lipid transport at contact sites between the ER and other organelles [6–10]. Among the twelve OSBP/ORP family members, OSBP, ORP1, ORP2 and ORP4 transfer cholesterol in a phosphatidylinositol 4-phosphate (PI4P)- and/or phosphatidylinositol 4, 5-bisphosphate (PI(4,5)P2)-dependent manner [8, 11]. Previously, we demonstrated that ORP2 functions to deliver cholesterol from internal organelles such as lysosomes to the PM in cultured cell lines [11]. However, the roles of ORP2 in physiological and pathological contexts have not been extensively examined.

Accumulation of free cholesterol and cholesterol crystals in hepatocytes is strongly associated with human Metabolic Dysfunction-Associated Steatohepatitis (MASH) [12–14]. Hepatocyte free cholesterol concentration is determined by several well-known pathways that include endogenous synthesis, uptake of cholesterol-containing plasma lipoproteins, cholesterol excretion into bile and conversion of cholesterol into bile acids [15]. In addition, free cholesterol can be esterified into hydrophobic cholesterol esters for storage within cytoplasmic lipid droplets (LDs) and for secretion within ER luminal lipoproteins [16]. Surprisingly however, gene expression studies failed to detect significant changes in those well-established pathways regulating cholesterol homeostasis in hepatocytes from MASH patients [12]. Therefore, additional factors/pathways may control free cholesterol accumulation in hepatocytes during the development of MASH.

The expression of ORP2 was reported to be significantly reduced in mouse liver after high-fat diet (HFD) treatment [17]. To examine the *in vivo* function of ORP2, we generated a liver-specific knock out mouse line of ORP2 (ORP2-LKO) in this study. We observed significant accumulation of free cholesterol and cholesterol crystals in the liver of ORP2-LKO mice fed HFD. Our results thus identify a novel pathway controlling the distribution of free cholesterol during MASH development.

## Results

### Generation and Characterization of ORP2-LKO Mice

A database search revealed that the transcription level of *Osbpl2/ORP2* was significantly decreased in mouse liver after high-fat diet (HFD) treatment compared with the normal chow diet (GSE 204986) [17](Fig 1A). Consistently, the protein level of ORP2 were significantly down-regulated in the high-fat-induced liver compared to normal diet as confirmed by Western blot (Fig 1B and 1C). To explore if and how *Osbpl2/ORP2* may regulate the physiological function of liver, we crossed *Osbpl2*^f/f^ mice with Alb-Cre mice and generated liver-specific *Osbpl2/ORP2* knock out mice, herein designated as ORP2-LKO mice (Fig 1D). The mRNA expression level of *Osbpl2* was dramatically and significantly decreased only in the liver (Fig. 1E). As shown in Fig 1F, the protein level of ORP2 was almost completely depleted in liver.

**Figure 1.**
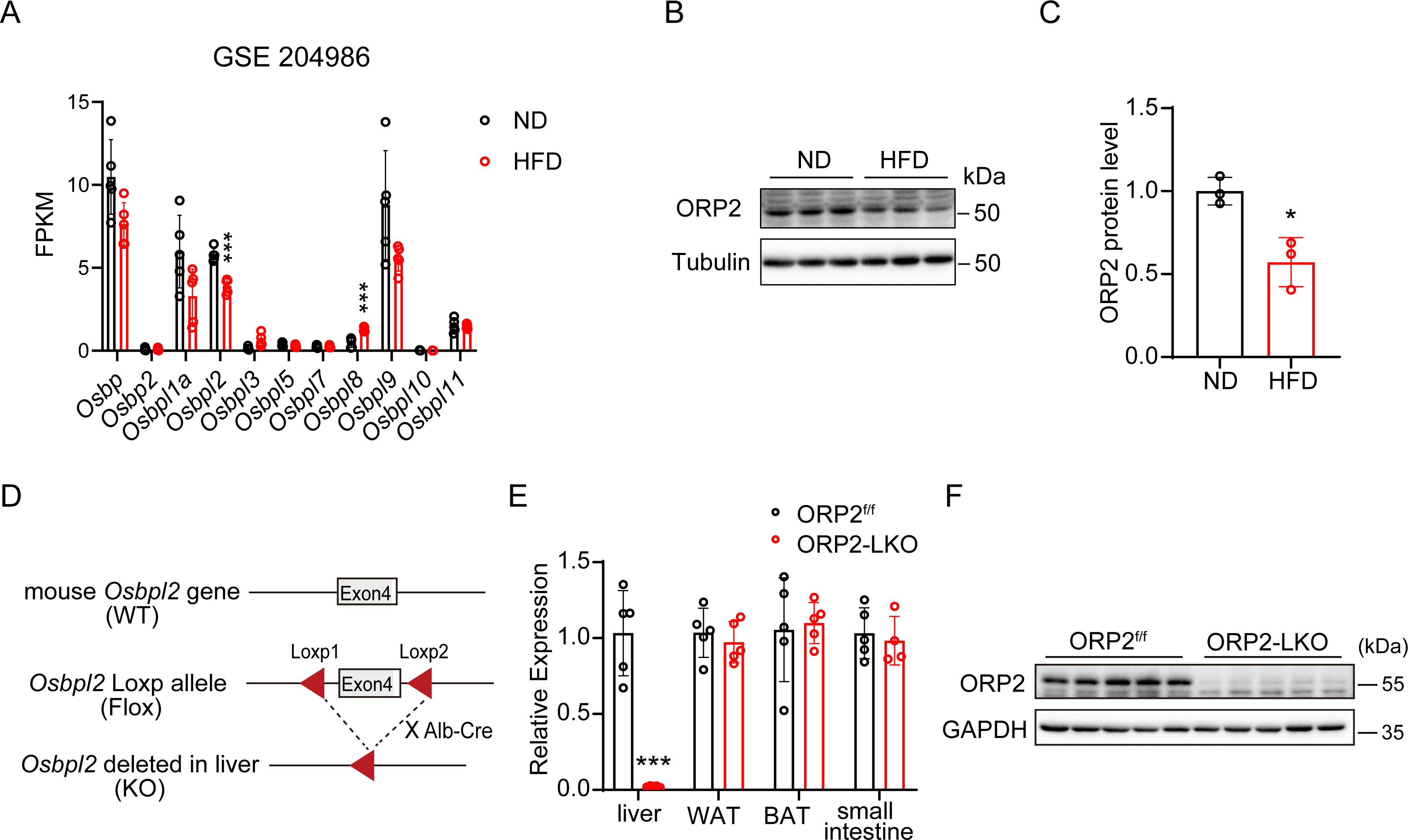
Association of ORP2 and fatty liver and construction of the ORP2-LKO mice. A. The FPKM (Fragments Per Kilobase of transcript per Million mapped reads) of ORP family members in liver of mice with the treatment of high fat diet (HFD) compared with normal diet (ND) from database GSE204986. B. Western blot analysis of ORP2 protein levels in liver of normal diet mice and high fat diet mice. C. Quantification of ORP2 protein level in Fig. 1B (n=3). D. Schematic diagram of ORP2 liver specific knockout mice construction. E. RT-qPCR analysis of relative mRNA expression of ORP2 in tissue of ORP2^f/f^ and ORP2-LKO mice (n=4-5). F. Western blot analysis of ORP2 protein levels in liver of ORP2^f/f^ and ORP2-LKO mice. Data are presented as mean ± SD. Statistical significance was determined using a two-tailed Student’s t-test. *P < 0.05, **P < 0.01, ***P < 0.001.

To determine the effect of liver ORP2 deficiency on whole body physiology, we first measured the body and organ weight of 9-month-old mice fed chow diet. No obvious differences in body weight, organ weight and fat weight were observed between ORP2-LKO and control mice in the rearing period (Fig S1A-S1C). The plasma level of TG (triglyceride), TC (total cholesterol), ALT (alanine aminotransferase) and AST (aspartate aminotransferase) did not display any notable differences between ORP2-LKO and floxed mice (Fig S1D-S1F). No obvious differences in lipid accumulation or inflammation in liver were detected between ORP2-LKO mice and *ORP2^f/f^* mice according to liver weight ratio (Fig S1G), the TG (Fig S1H) and TC (Fig S1I) content in liver, as well as H&E, Oil red O, F4/80 and Sirius red staining analyses (Fig S1J). These results suggest that the absence of liver ORP2 did not have a strong metabolic impact on whole body and liver physiology under normal chow diet.

### The absence of liver ORP2 exacerbated diet-induced obesity

To examine if liver ORP2 may play any role in HFD-induced metabolic abnormalities, 8-week-old ORP2-LKO and ORP2^f/f^ mice were fed with HFD for 8 weeks. The weight gain of both ORP2-LKO mice and the control litter mates remain comparable with each other during the first few weeks of HFD treatment, whereas ORP2-LKO mice gained significantly more weight after seven weeks on HFD (Fig 2A). The body composition of mice measured by nuclear magnetic resonance showed that the fat mass of ORP2-LKO was significantly higher than control mice (Fig 2B), whereas the lean mass ORP2-LKO of mice was not changed (Fig 2C). The spleen of the ORP2-LKO mice showed a moderate but significant reduction in weight, but no weight differences were detected in heart and kidney between WT and ORP2-LKO mice (Fig. 2D). Among fat depots, the weight of subcutaneous WAT (sWAT) of ORP2-LKO mice increased slightly (Fig 2E), whereas the weight of the gonadal WAT (gWAT), brown adipose tissue (BAT) showed little change (Fig 2E). Interestingly, histology analyses showed that lipid droplets expansion expanded in the SWAT (Fig 2F and 2G) and GWAT (Fig 2H and 2I) of ORP2-LKO mice fed HFD. To further characterize these mice, we subjected them to metabolic cage analysis. The respiratory exchange ratio (Fig 2J), oxygen consumption (Fig S2A) and carbon dioxide production (Fig S2B) of ORP2-LKO mice remain similar to those of ORP2^f/f^ mice. After loading with high fat diet for 8 weeks, no significant difference was detected in glucose and insulin tolerance tests between ORP2-LKO and WT mice (Fig S2C-S2D). Overall, these results indicated that the deletion of liver ORP2 had moderate impact on whole body metabolism.

**Figure 2.**
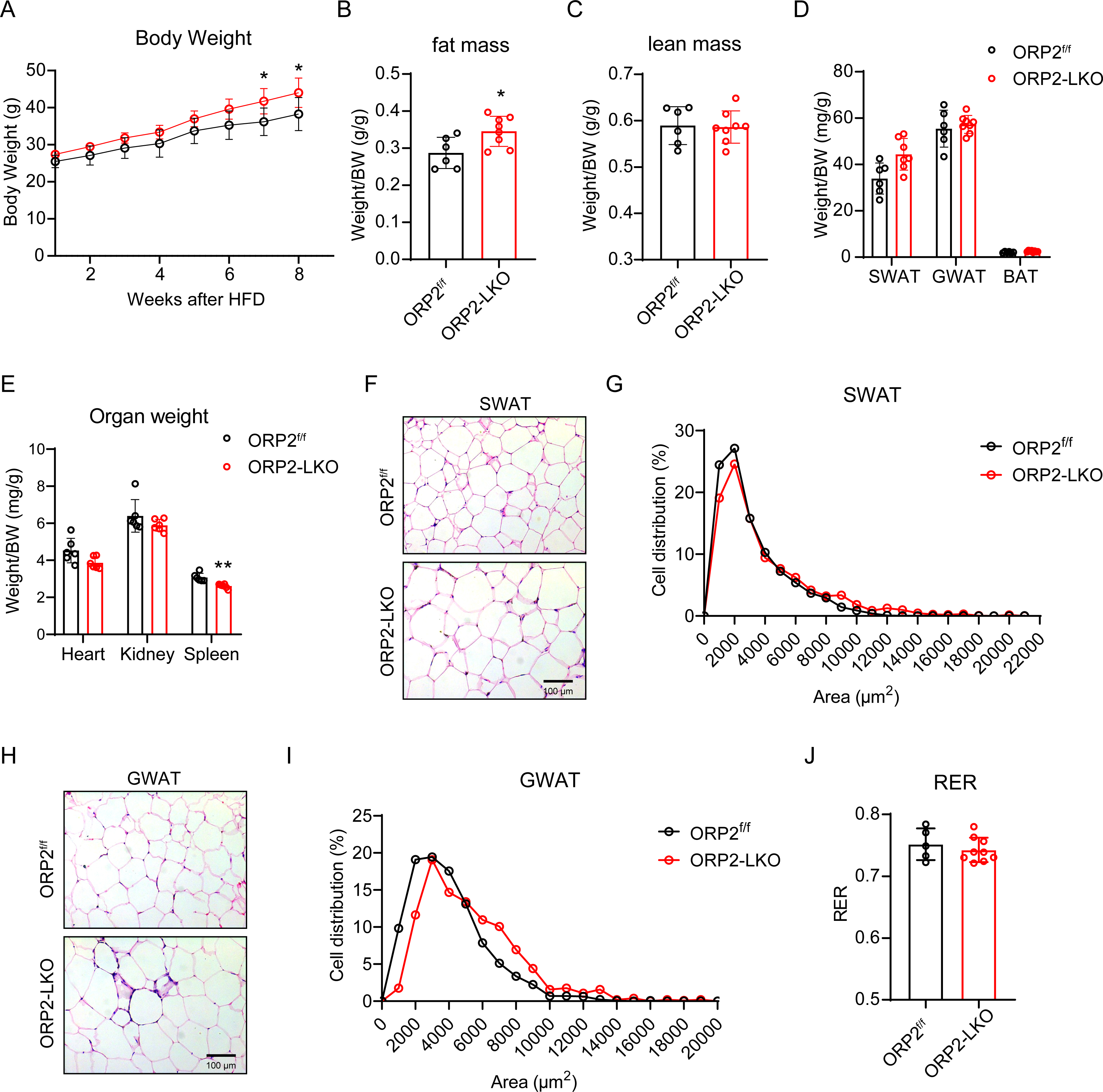
The absence of liver ORP2 exacerbated diet-induced obesity. A. Growth curve of ORP2^f/f^ mice and ORP2-LKO mice during HFD feeding (n=6-8). B and C. Fat mass and lean mass of ORP2^f/f^ mice and ORP2-LKO mice measured by nuclear magnetic resonance (NMR) analyzer after 8 weeks HFD feeding (n=6-8). D. Fat weight of ORP2^f/f^ mice and ORP2-LKO mice (n=6-8). E. Organ weight of ORP2^f/f^ mice and ORP2-LKO mice (n=5-6). F and G. H&E stain of Subcutaneous fat (SWAT) and quantification of adipocyte size of SWAT in ORP2^f/f^ mice and ORP2-LKO mice. H and I. H&E stain of Gonadal fat (GWAT) and quantification of adipocyte size of GWAT in ORP2^f/f^ mice and ORP2-LKO mice. J. Respiratory exchange ratios of mice subjected to metabolic cage analysis (n=5-9). Data are presented as mean ± SD. Statistical significance was determined using two-way ANOVA was performed, followed by Sidak’s multiple comparison test for Figure 2A and two-tailed Student’s t-test for Figure 2B and 2D. *P < 0.05, **P < 0.01, ***P < 0.001.

### The deficiency of liver ORP2 exacerbated HFD-induced liver steatosis

The ORP2-LKO mice showed higher levels of plasma total cholesterol (TC) (Fig 3A) and triglycerides (TG) (Fig 3B), as well as plasma alanine aminotransferase (ALT) (Fig 3C) and aspartate aminotransferase (AST), indicating liver damage (Fig 3D). Furthermore, the liver of ORP2-LKO was moderately but significantly heavier than control mice (Fig 3E). As revealed by H&E analysis and Oil red O staining, the ORP2-LKO livers developed severe hepatic steatosis, as marked by vacuolated and lipid-laden hepatocytes (Fig 3F). Consistently, the TG (Fig 3G) and TC (Fig 3H) content in the liver of ORP2-LKO mice were significantly elevated as compared with floxed control mice. Interestingly, the level of free cholesterol displayed a significant increase, whereas cholesterol esters content remained similar between ORP2-LKO liver and normal liver (Fig 3H). Consistent with increased free cholesterol, more cholesterol crystals were observed with a polarized light microscope in ORP2-LKO mice (Fig 3I). Together, these results suggest that ORP2 functions to maintain normal lipid, especially cholesterol metabolism in the liver during HFD-induced MASH development.

**Figure 3.**
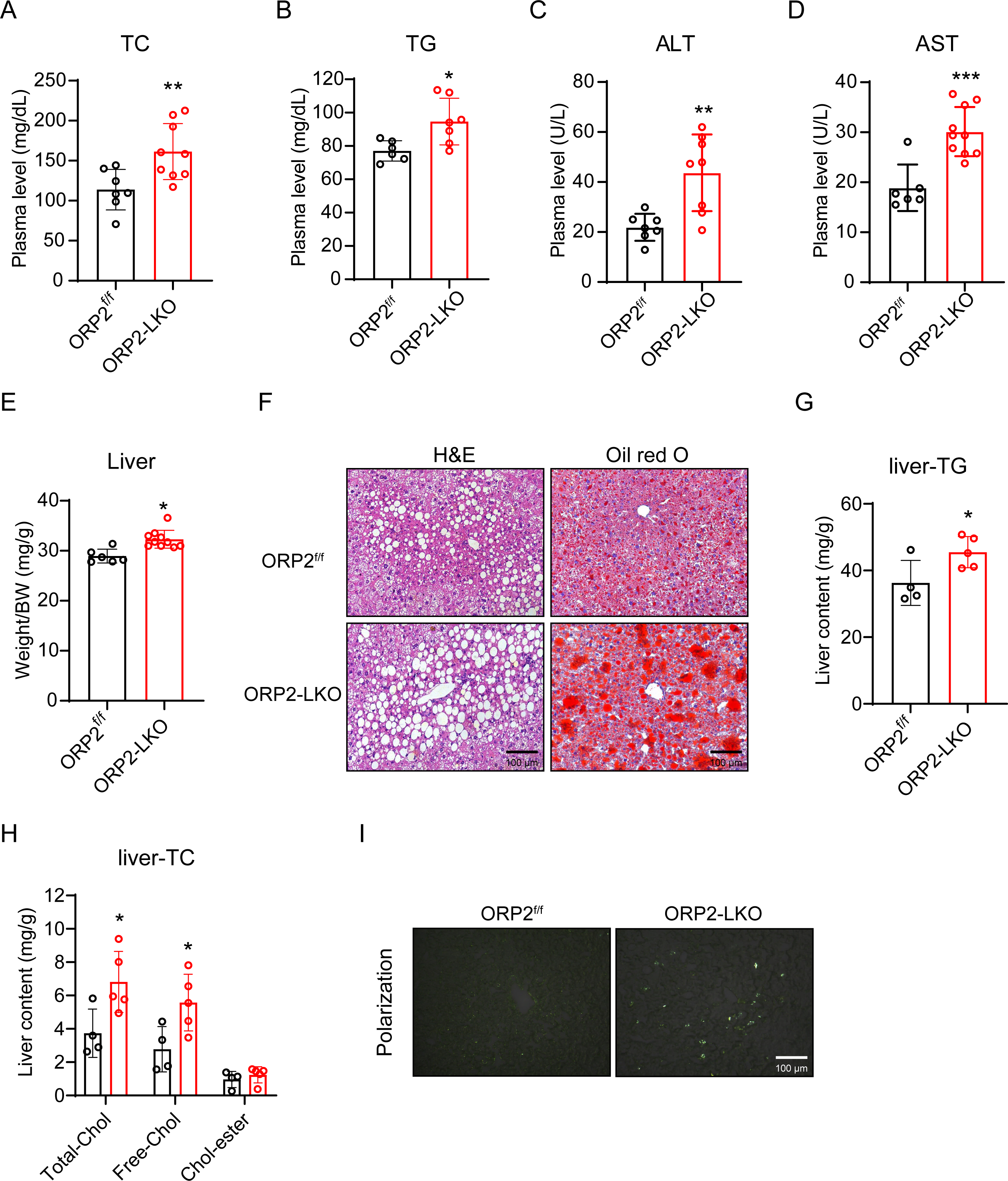
ORP2 deficiency exacerbated hepatic steatosis and free cholesterol accumulation. A, B, C and D. Plasma level of total cholesterol (TC), triglyceride (TG), ALT and AST in ORP2^f/f^ mice and ORP2-LKO mice after 8 weeks of HFD feeding (n=6-10). E. The ratio of liver weight to body weight of ORP2^f/f^ mice and ORP2-LKO mice (n=6-10). F. The H&E and Oil red O staining in the liver of ORP2^f/f^ mice and ORP2-LKO mice. Scale bar 100 μm. G. The content of TG in liver of ORP2^f/f^ mice and ORP2-LKO mice (n=4-5). H. The content of total-cholesterol (Total-Chol), free-cholesterol (Free-Chol) and cholesterol-ester (Chol-ester) in liver of ORP2^f/f^ mice and ORP2-LKO mice (n=4-5). I. Representative polarization microscopy images of liver sections from ORP2^f/f^ mice and ORP2-LKO mice. Scale bar 100 μm. Data are presented as mean ± SD. Statistical significance was determined using a two-tailed Student’s t-test. *P < 0.05, **P < 0.01, ***P < 0.001.

### Deletion of ORP2 exacerbates HFD-induced hepatic inflammation and fibrosis

Many studies have reported that increased lipid or cholesterol accumulation in the liver can trigger inflammation and fibrosis [13, 18]. In order to explore the role of ORP2 in the development of inflammation in the liver on HFD, we performed a series of experiments to examine inflammation in WT and ORP2-deficient liver. Firstly, the staining of F4/80 in liver was significantly stronger in ORP2-LKO mice than in the controls, with more crown like structures (CLS) (Fig 4A and 4B). Moreover, the expression level of inflammation-related genes such as TNFα was significantly increased in liver after ORP2 deletion (Fig 4C). Furthermore, the population of Ly6C^high^ pro-inflammatory macrophages was significantly increased in the liver of ORP2-LKO mice, as detected by flow cytometry (Fig 4D and 4E). In addition, elevated fibrosis is evident in the liver of ORP2-LKO mice as revealed by Sirius red staining (Fig 4F) and the mRNA level of *Col3a1* (Fig 4G). Together, these data suggest that HFD-induced inflammation and fibrosis of liver were exacerbated by the loss of liver ORP2.

**Figure 4.**
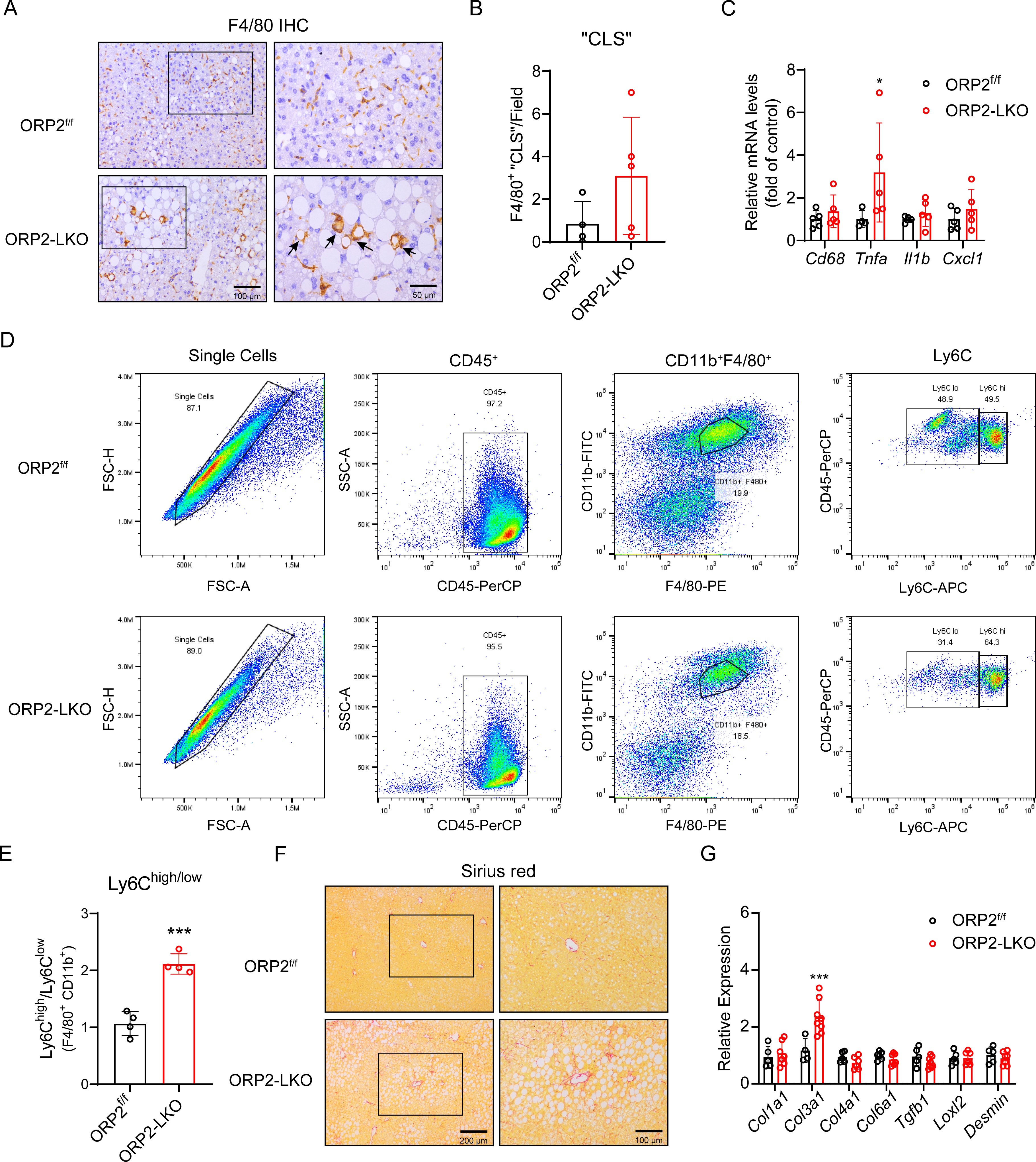
Severe hepatic inflammation and fibrosis in ORP2-LKO mice. A and B. Representative immunohistochemical staining images of F4/80 (A) and quantification of F4/80 positive crown like structure (B) in liver section from ORP2^f/f^ and ORP2-LKO mice fed a high-fat diet for 8 weeks (n = 4 - 5). Scale Bar: 100 or 50 µm. C. RT-qPCR analysis of relative mRNA expression of inflammation genes in the liver from ORP2^f/f^ and ORP2-LKO mice fed a high-fat diet for 8 weeks (n = 5-7). D and E. Representative fluorescence-activated cell sorting (FACS) analysis and quantification of Ly6C^high^/Ly6C^low^ macrophages gated on the F4/80^+^CD11b^+^ population within the CD45^+^ cells from ORP2^f/f^ and ORP2-LKO mice fed a high-fat diet for 8 weeks (n = 4). F. Representative Sirius red staining images in liver sections from ORP2^f/f^ and ORP2-LKO mice fed a high-fat diet for 8 weeks (n = 4 - 5). Scale Bar: 200 or 100 µm. G. RT-qPCR analysis of relative mRNA expression of fibrosis genes in the liver from ORP2^f/f^ and ORP2-LKO mice fed a high-fat diet for 2 months (n = 5-7). Data are presented as mean ± SD. Statistical significance was determined using a two-tailed Student’s t-test. *P < 0.05, **P < 0.01, ***P < 0.001.

### ORP2 is required for normal bile acid metabolism in liver

To gain additional insights into how ORP2 may control hepatic metabolism, we performed metabolomics and RNA-sequencing analyses on liver tissues from ORP2^f/f^ and ORP2-LKO mice fed with HFD. Metabolomics analyses results showed that the levels of 46 metabolites were increased and those of 393 metabolites decreased (Fig 5A). Among them, differences in bile acid metabolites between the two groups were particularly significant (Fig 5B). Especially, taurocholic acid, a key component of bile acids, was significantly increased in ORP2-deficient liver (Fig 5C). In the bile, bile acids, phosphatidylcholine (PC), and cholesterol form mixed micelles, which are stored in the gallbladder [19]. Metabolomics analyses also revealed that the levels of two species of PC (PC(20:3(8Z,11Z,14Z)-2OH(5,6)/18:2(9Z,12Z)) and PC(18:2(9Z,12Z)/17:1(9Z))) increased markedly in ORP2-LKO mice compared with control mice (Fig 5D-5E). Kyoto Encyclopedia of Genes and Genomes (KEGG) pathway enrichment analysis from metabolomics results revealed a strong enrichment of genes involved in bile secretion and cholesterol metabolism (Fig 5F). These data suggest that ORP2 may regulate bile acid metabolism through its role in maintaining cellular cholesterol homeostasis.

**Figure 5.**
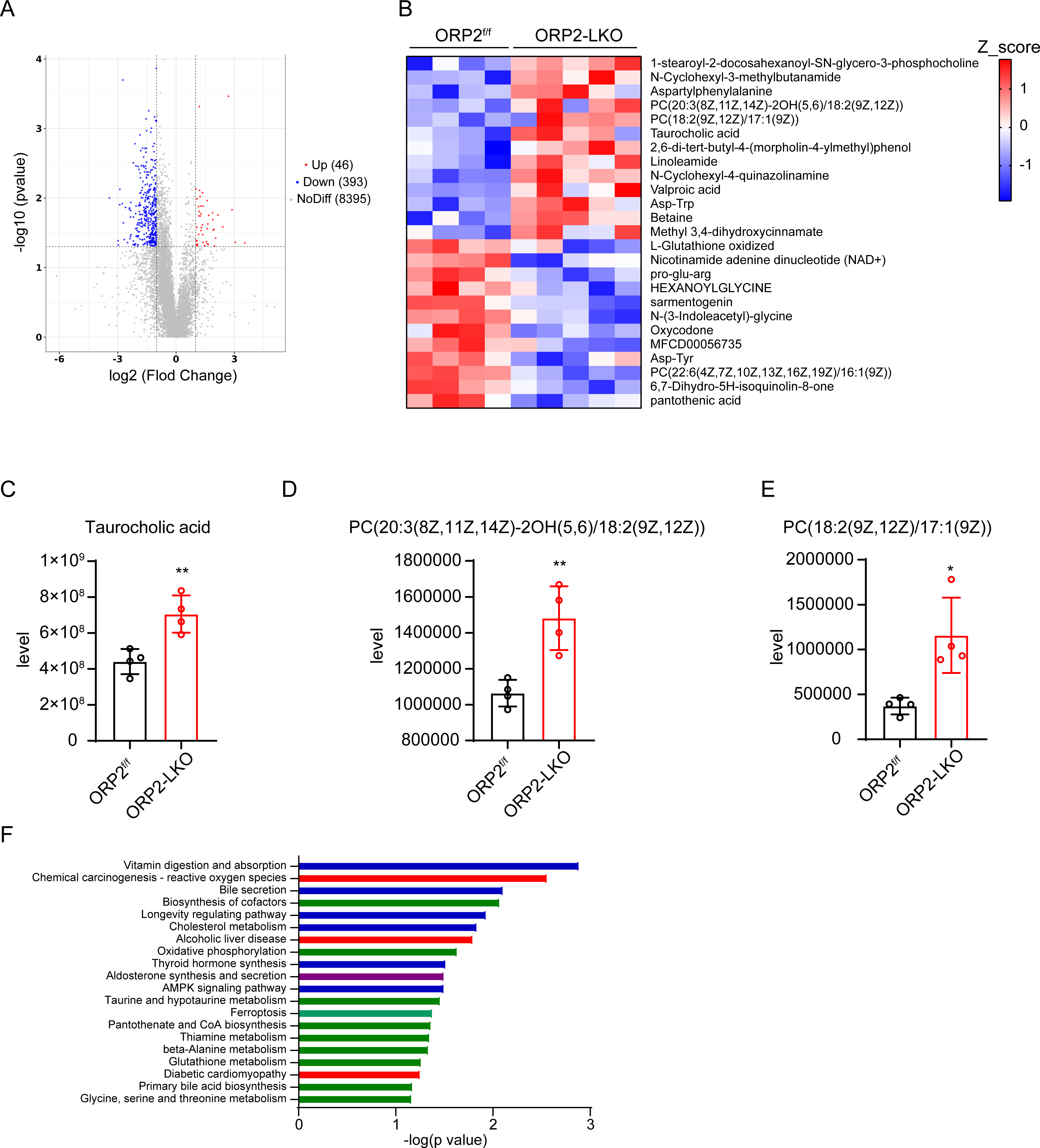
Metabolomics analysis of liver in ORP2^f/f^ and ORP2-LKO mice on high fat diet. A. Volcano map of metabolomics analysis of liver in ORP2^f/f^ and ORP2-LKO mice. 46 metabolites were upregulated, 393 metabolites were down-regulated and 8395 metabolites had no difference in ORP2-LKO mice. B. Heat map of metabolomics of liver in ORP2^f/f^ and ORP2-LKO mice. C, D and E. The level of taurocholic acid, PC (20:3/18:2) and PC (18:2/17:1) in liver in ORP2^f/f^ and ORP2-LKO mice (n = 4). F. Metabolomics KEGG pathway enrichment analysis in liver in ORP2^f/f^ and ORP2-LKO mice. Data are presented as mean ± SD. Statistical significance was determined using a two-tailed Student’s t-test. *P < 0.05, **P < 0.01, ***P < 0.001.

To further evaluate the effect of ORP2 on liver metabolism, we extracted mRNA from the liver of ORP2-LKO and their littermate control and performed RNA sequencing (RNA-seq). Among the 16,819 genes analyzed, 57 genes were upregulated in ORP2-LKO mice, including the bile acid related genes, such as *Cyp7a1* and *Onecut1* (Fig 6A). Heat map of RNA-sequencing analysis showed that the expression of primary bile acid biosynthesis (Gene cluster 3 (G-C3)) related genes was upregulated significantly (Fig 6B). KEGG pathway enrichment analysis from RNA-seq revealed a strong enrichment of genes involved in steroid biosynthesis, such as steroid hormone biosynthesis (Fig 6C). To confirm these results, we examined the expression of some key genes responsible for bile acid synthesis, export, uptake, efflux and signaling pathway by qPCR. The expression of *Cyp7a1* (bile acid synthesis) and *Onecut1* (bile acid signaling) were dramatically increased. By contrast, the expression of *Abcg5* and *Abcg8,* critical to cholesterol efflux, was significantly decreased in ORP2-deficeincy liver (Fig 6D). Taken together, ORP2 deficiency led to disturbed bile acid metabolism and increased bile acid accumulation in liver.

**Figure 6.**
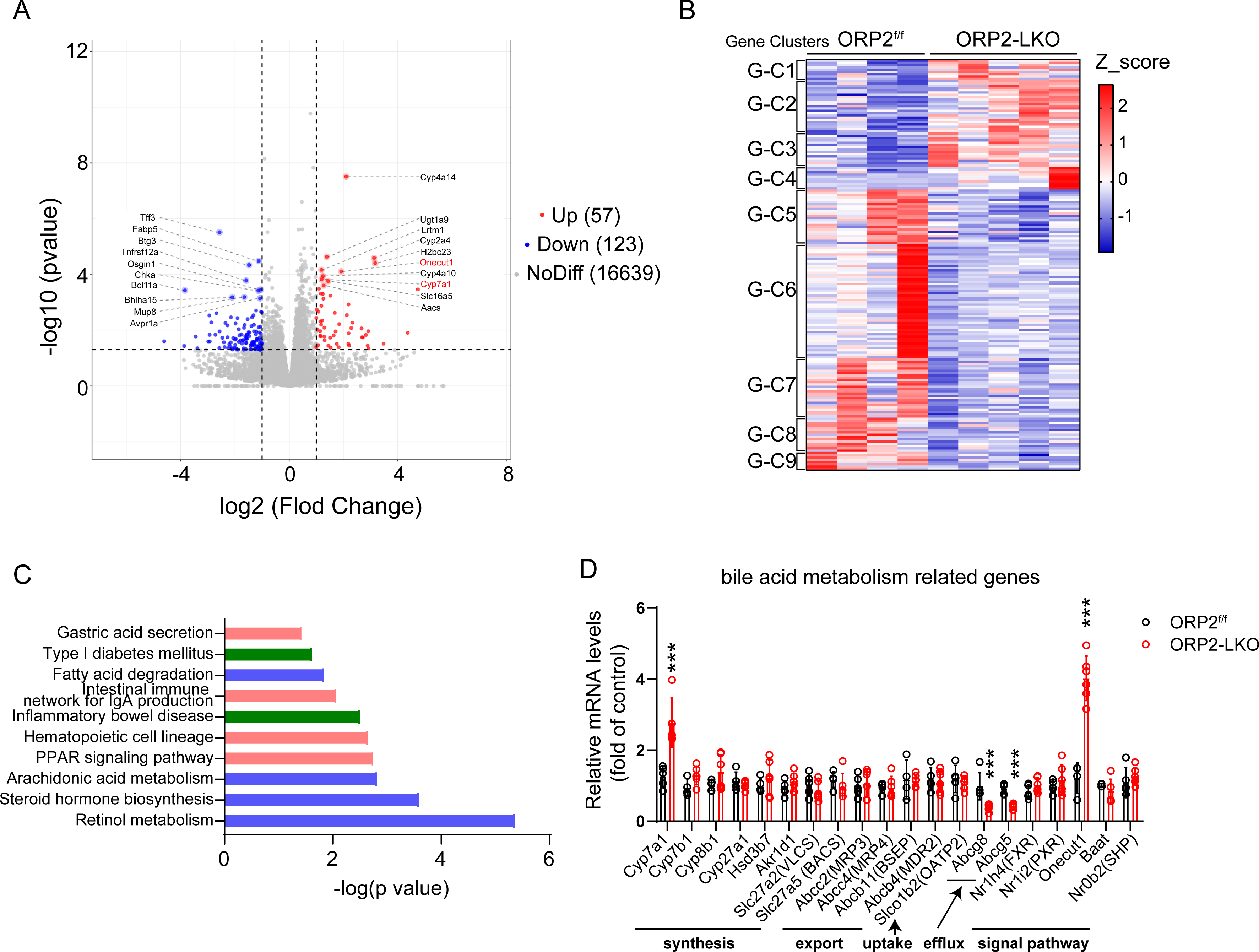
RNA-sequencing analysis of livers from ORP2^f/f^ and ORP2-LKO mice on high fat diet. A. Volcano map of RNA-sequencing results of liver in ORP2^f/f^ and ORP2-LKO mice. 57 genes were up-regulated, 123 genes were down-regulated and 16639 genes had no difference in ORP2-LKO mice as compared with the control mice. B. Heat map of RNA-sequencing analysis of liver in ORP2^f/f^ and ORP2-LKO mice. Gene cluster1 (G-C1): Histidine metabolism, G-C2: Retinol metabolism (fatty acid degradation), G-C3: Primary bile acid biosynthesis, G-C4: Arachidonic acid metabolism, G-C5: Linoleic acid metabolism, G-C6: Cytokine-cytokine receptor interaction (inflammatory bowel disease), G-C7: Cytokine-cytokine receptor interaction (hypertrophic cardiomyopathy), G-C8: Maturity onset diabetes of the young, G-C9: Linolenic acid metabolism. C. KEGG pathway enrichment analysis of RNA-sequencing results of liver from ORP2^f/f^ and ORP2-LKO mice. D. Relative mRNA levels of bile acid metabolism related genes analyzed by RT-qPCR. Data are presented as mean ± SD. Statistical significance was determined using a two-tailed Student’s t-test. *P < 0.05, **P < 0.01, ***P < 0.001.

## Discussion

Increased free cholesterol content in the liver is one of the hallmarks of human MASH [20]. Moreover, free cholesterol crystals were specifically found in patients with MASH, but not in patients with simple steatosis [12]. Thus, crystals of free cholesterol may causally contribute to the development of MASH. For instance, free cholesterol may promote MASH through the stabilization and activation of TAZ [21]. Although the important role of free cholesterol in the development of MASH has been well recognized, much is to be learned about the genetic factors and molecular mechanisms that govern the hepatic accumulation of free cholesterol during MASH development.

Previously, we and others demonstrated that the primary function of ORP2 is to deliver free cholesterol to the plasma membrane from endosomal compartments [11, 22, 23]. ORP2 deficiency caused free cholesterol to accumulate in the endolysomal membranes in cultured cell lines. In the current study, we show that upon HFD treatment, ORP2 deficiency caused significant accumulation of free cholesterol in the liver, accompanied by increased inflammation and liver damage. It is conceivable that ORP2 deficiency in hepatocytes may cause free cholesterol to accumulate in the endolysomal membranes. The accumulated free cholesterol may then move directly to lipid droplets for storage, as suggested [12]. Upon HFD feeding, more and more cholesterol enters the endolysosomal system of hepatocytes, accumulates on LD surface, and ultimately exceeds the capacity of LD surface phospholipids to accommodate cholesterol. Consequently, free cholesterol may then precipitate and form crystals next to LDs. Since less cholesterol can reach the plasma membrane of ORP2-deficient hepatocytes, cholesterol efflux may be reduced as suggested by the decrease in ABCG5/8 expression, further exacerbating intracellular cholesterol accumulation. The increase in CYP7A1 expression is likely a response to increased cellular free cholesterol. Finally, it should be noted that the HFD used in this study does not usually cause hepatic accumulation of free cholesterol because it is not supplemented with cholesterol [21]. Therefore, our data suggest that the loss of ORP2 in hepatocytes significantly accelerated free cholesterol accumulation and crystallization.

There are a number of limitations in this study. For instance, we cannot determine if endolysomal cholesterol is increased in ORP2-deficient hepatocytes. It is also somewhat confusing why cholesterol esters did not accumulate in ORP2-deficient hepatocytes. Perhaps the excessive free cholesterol is not efficiently esterified or the cholesterol esters are more efficiently incorporated into lipoproteins and secreted in ORP2-deficient hepatocytes. The increased plasma total cholesterol supports the latter possibility. Moreover, we cannot explain why only taurocholic acid, but not other bile acids, is selectively upregulated in ORP2-deficient liver. Future studies will address the underlying molecular mechanisms of the observations reported here.

In summary, we have generated a mouse model to examine ORP2 function *in vivo*, and our results are consistent with the known biochemical function of ORP2. Most importantly, we identify ORP2 as a novel molecule that regulates the accumulation of hepatic free cholesterol during the development of MASH. Our results therefore indicate that proper intracellular cholesterol trafficking may represent an important regulatory node to prevent free cholesterol accumulation during MASH.

## Methods

### Mice

The Osbpl2^f/f^ mice were generated by Cyagen Biosciences (Guangzhou, CHN), then bred with Alb-cre transgenic mice to generate liver specific knockout mice. The high fat diet contained 60% calories from fat, 20% calories from carbohydrates, and 20% calories from protein. Mice were housed in the animal facility with 12-h light/12-h dark cycles, the temperature at 22°C-23°C and 20-60% humidity with free access to diet and water. Male mice are used in the studies. All animal procedures were approved by the Laboratory Animal Ethical and Welfare Committee of Hebei Medical University, and the Institutional Ethics Committee of Fudan University and Nanjing University, under a permit of animal use in the Center of Experimental Animal at Shanghai and Nanjing. The permit followed the Experimental Animal Regulations set by the National Science and Technology Commission, China.

Body Composition Measurement: The minispec LF50 nuclear magnetic resonance (NMR) analyzer (Bruker) was used to measure the body composition of mice according to the manufacturer’s instructions.

Glucose tolerance test and insulin tolerance test: For the glucose tolerance test, mice were fasted for 14-16 h and then i.p. given glucose (2 g of glucose/kg). Blood glucose was measured at 0, 15, 30, 60, and 120 min after injection of glucose. For insulin tolerance test: mice were fasted for 6 h before experiment. Insulin (0.7 U/kg) was administered i.p., and blood glucose was measured at 0, 15, 30 and 60 min after injection.

Indirect calorimetry measurement: Metabolic cage analysis was performed in Comprehensive Laboratory Animal Monitoring System (CLAMS) (Columbus Instruments). Mice were acclimatized to the metabolic cages for 24 h. Mice were then monitored for 2 days on food intake, body weight, oxygen consumption, carbon dioxide production, and locomotor activities. The RER (respiratory exchange ratio) is calculated as: RER=VCO2/VO2

Detection of serological indicators: LabAssay^TM^ Triglyceride (Wako, 290-63701), LabAssay^TM^ Total cholesterol (Wako, 635-50981), Aspartate aminotransferase Assay Kit (AST) (Nanjing Jiancheng, C010-2-1), Alanine aminotransferase Assay Kit (ALT, Nanjing Jiancheng, C009-2-1). RNA-seq and Metabolomics in liver were analyzed by Personalbio.

### RNA extraction and RT-qPCR analysis

Total RNA from liver were extracted with Trizol regent (15596026CN, ThermoFisher). Complementary DNA was synthesized using the reverse transcription kit (11151ES10, Yeasen). RT-qPCR was performed with Hieff® qPCR SYBR Green Master Mix (11202ES03, Yeasen). The primers used for RT-qPCR were listed in Table 1.

**Table 1:**
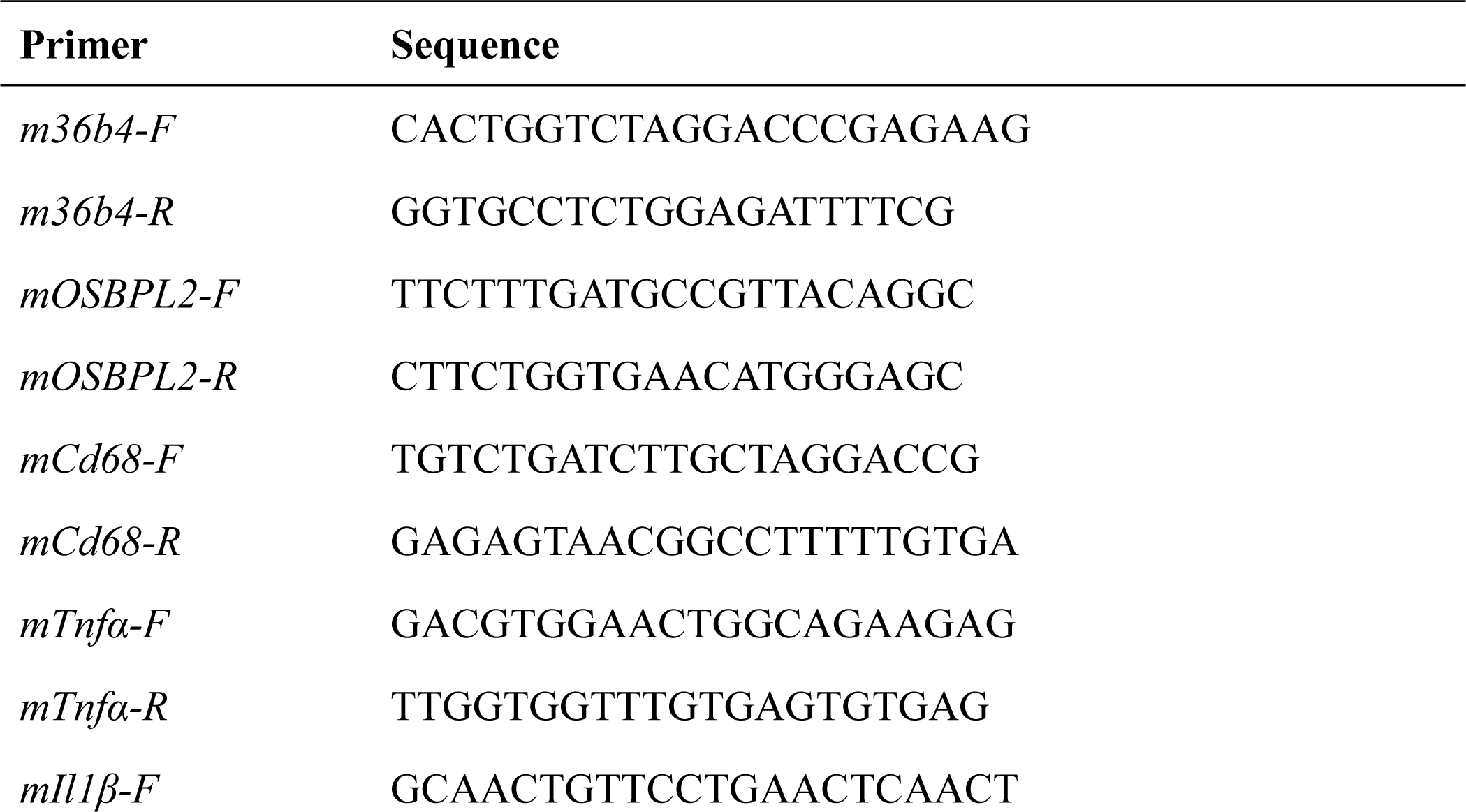
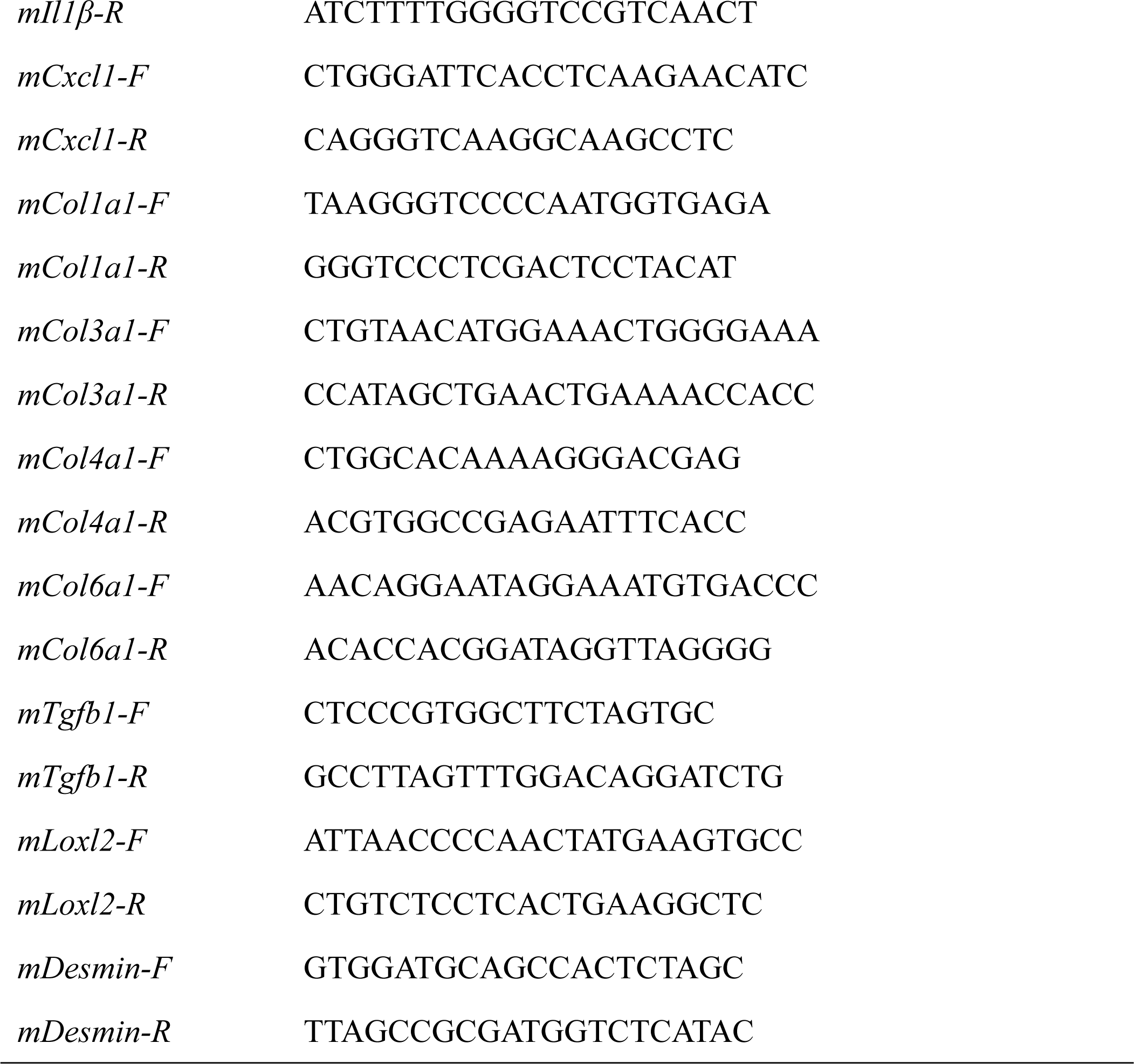
List of the primers used in this study.

### Western blot

Approximately 100 mg of tissues were resuspended and homogenized with lysis solution (50 mM Tris, 150 mM NaCl, 1% NP-40, 0.5% sodium deoxycholate, 5 mM EDTA containing protease inhibitors (P1005, Beyotime)) and lysed for 15 min on ice, then supernatant was collected after centrifuged 12000 rpm for 15 min at 4℃. Protein concentrations were quantified using BCA protein assay kit (P0012, Beyotime). Equal amounts of protein were loaded onto SDS-PAGE and transferred to PVDF membranes. After blocked with 5% milk for 1 h at room temperature, primary antibodies were diluted in primary antibody dilution buffer (P0023A, Beyotime) and incubated with membranes overnight at 4℃. Membranes were washed three times with TBST at room temperature for 6 min per wash. Secondary antibodies (HRP-conjugated goat anti– mouse IgG (115-035-003, Jackson), HRP-conjugated goat anti–rabbit IgG (111-035-003, Jackson)) were diluted in 5% milk and incubated with membranes for 1 h at room temperature. Western blots were imaged using the Bio-Rad system. Quantification of band intensities was performed using ImageJ (NIH). The following primary antibodies were used for western blot: Rabbit polyclonal anti-OSBPL2 (14751-1-AP, Proteintech), Mouse monoclonal-β-Tubulin (AC021, Abclonal), Mouse monoclonal GAPDH (AC002, Abclonal).

### Liver Lipid Extraction and Assay

Approximately 100 mg of frozen liver tissue was homogenized in 1 mL of cold PBS using a homogenizer. Lipids were extracted with chloroform/methanol (2:1, v/v), then dried under nitrogen and re-dissolved in 0.5 mL of 3% Triton X-100.

Liver triglyceride content and total cholesterol content were measured using assay kits from Bio Sino (Beijing, China), following the manufacturer’s instructions. Liver cholesteryl ester content was calculated as the difference between total cholesterol and free cholesterol levels. The assays for liver total cholesterol and free cholesterol content were performed using kits from Applygen (Beijing, China), also according to the manufacturer’s instructions.

### Histological Analysis

The liver and adipose tissues were fixed and embedded in paraffin. Sections were cut at a thickness of 7 μm and stained with hematoxylin and eosin (H&E). Additionally, after fixation, the liver was embedded in OCT (Sakura Finetek, USA) and 7 μm thick sections were obtained by cryosectioning. Lipid deposition in these cryosections was assessed using Oil-Red O staining. Cholesterol crystals in frozen sections were observed with a polarized light microscope (SOPTO CX40P, Suzhou, China).

For paraffin-embedded sections, immunohistochemical analysis was performed using an F4/80 antibody (Servicebio, Wuhan, China) to evaluate macrophage infiltration in liver tissue. Sirius Red staining (Solarbio, Beijing, China) was used to assess liver fibrosis.

Adipocyte quantification was conducted using ImageJ software. For each experimental group, 5 mice were analyzed, with 5 fields of view at 200x magnification selected per mouse. The area of each adipocyte in these fields was measured and quantified.

### Flow Cytometry Analysis of Liver Ly6C^high^ and Ly6C^low^ Macrophages

Disaggregate approximately 500 mg of liver tissue mechanically to isolate non-parenchymal cells. Resuspend the cells in 0.5% BSA solution and incubate at room temperature for 15 minutes to block non-specific binding. Stain the cells with the following antibodies for 30 minutes: CD45-PerCP, F4/80-PE, CD11b-FITC, and Ly6C-APC (all from Tonbo Biosciences). Then analyze the stained cells using an Agilent flow cytometer (USA). First, gate on the CD45-positive cell population. Within the CD45-positive gate, further identify and select the CD11b and F4/80 double-positive macrophages. Then analyze the proportions of Ly6C high and Ly6C low cells within the CD11b and F4/80 double-positive macrophage subset.

### Statistical analysis

Data were presented as mean ± SD. Statistical analyses were performed using GraphPad Prism 8 software. The significance of differences between two groups was assessed using Student’s t-test. When comparing with more than two variables, two-way ANOVA was performed, followed by Sidak’s multiple comparison test. A P-value of <0.05 was considered statistically significant.

## ACKNOWLEDGEMENT

This work was supported by grants from the National Natural Science Foundation of China (92357308, 92357302, and 32270724 to F.J.C.), the National Key R&D Program of China (2018YFA0800301 to F.J.C.). M.G was supported by Natural Science Foundation of Hebei province (Grant No. H2022206462 and H2022206597). S.C. and H.Y.W. were supported by the Ministry of Science and Technology of China (Grant No. 2018YFA0801100). S.C. was also supported by the National Natural Science Foundation of China (Grant No. 32025019). H.Y. was supported by an Investigator Grant (2009852 from the National Health and Medical Research Council (NHMRC) of Australia, and by start-up funding from the University of Texas Health Science Center at Houston.

The authors declare no competing financial interests.

## Author contributions

J. Wu, Y. Zhao, L. Qiu, Y. Liang, Q. Chen, X. Wang, J. Gu performed experiments. All authors analyzed data. J. Wu, H. Wang, S. Chen, Y. Liu, X. Wu, F.J. Chen, M. Gao and H. Yang designed research. J. Wu, M. Gao and H. Yang wrote the manuscript with input from all other authors.

---

**Supplemental Figure 1.**
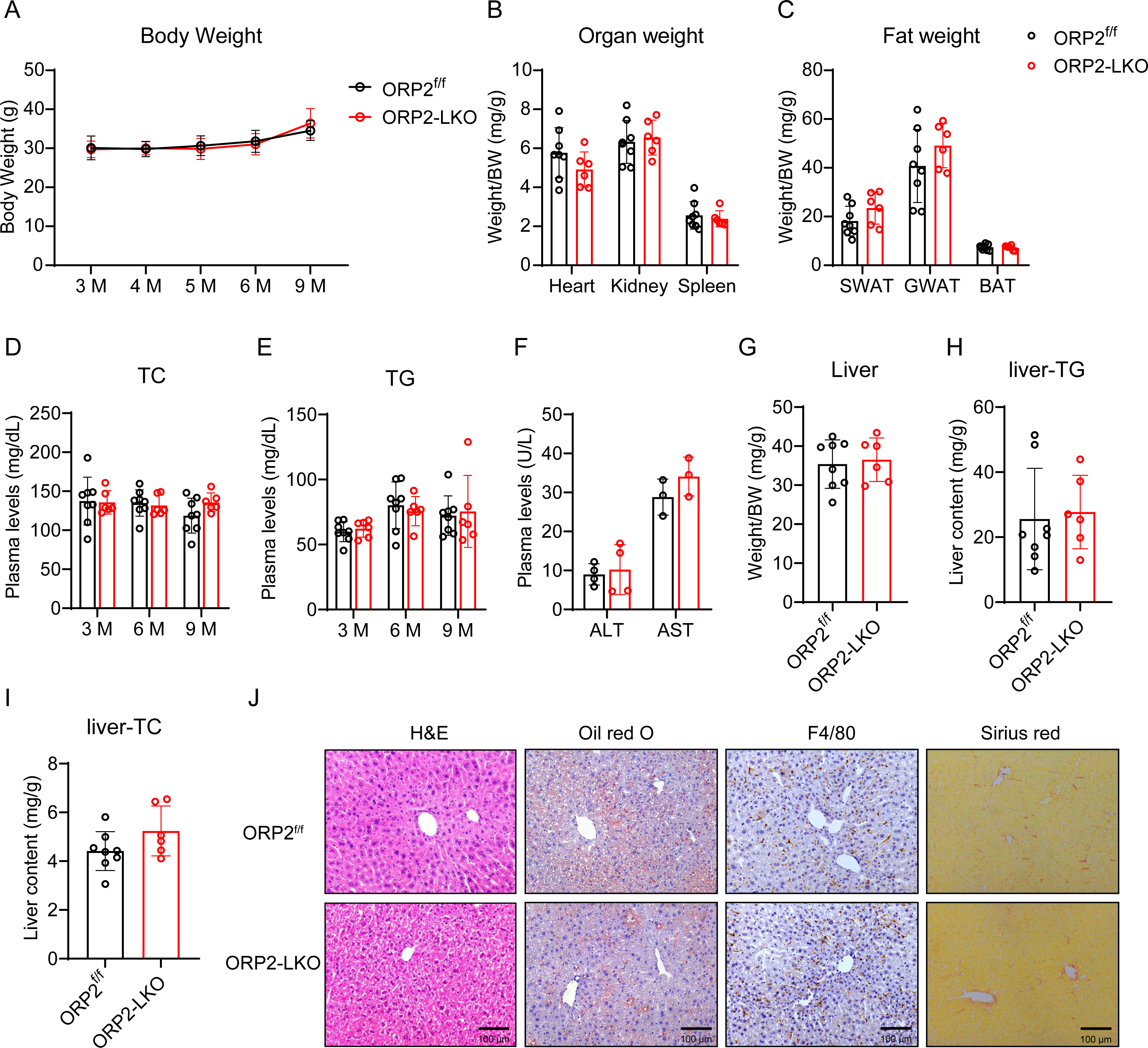
Metabolic characterization of ORP2 LKO mice on chow diet. A. Growth curve of ORP2^f/f^ mice and ORP2-LKO mice in normal diet (n=6-8). B. Organ weight of ORP2^f/f^ mice and ORP2-LKO mice in normal diet (n=6-8). C. Fat weight of ORP2^f/f^ mice and ORP2-LKO mice in normal diet (n=6-8). D and E. Plasma level of total cholesterol (TC) and triglyceride (TG) in ORP2^f/f^ mice and ORP2-LKO mice at 3, 6, and 9 months of age (n=6-8). F. Plasma level of ALT and AST in ORP2^f/f^ mice and ORP2-LKO mice at 9 months of age (n=6-8). G. The ratio of liver weight to body weight of ORP2^f/f^ mice and ORP2-LKO mice (n=6-8). H and I. The content of TG and TC in liver of ORP2^f/f^ mice and ORP2-LKO mice (n=6-8). J. The H&E stain and Oil red O stain F4/80 immunofluorescence stain and Sirius red stain of liver in ORP2^f/f^ mice and ORP2-LKO mice. Scale bar, 100 μm. Data are presented as mean ± SD. Statistical significance was determined using a two-tailed Student’s t-test. *P < 0.05, **P < 0.01, ***P < 0.001.

**Supplemental Figure 2.**
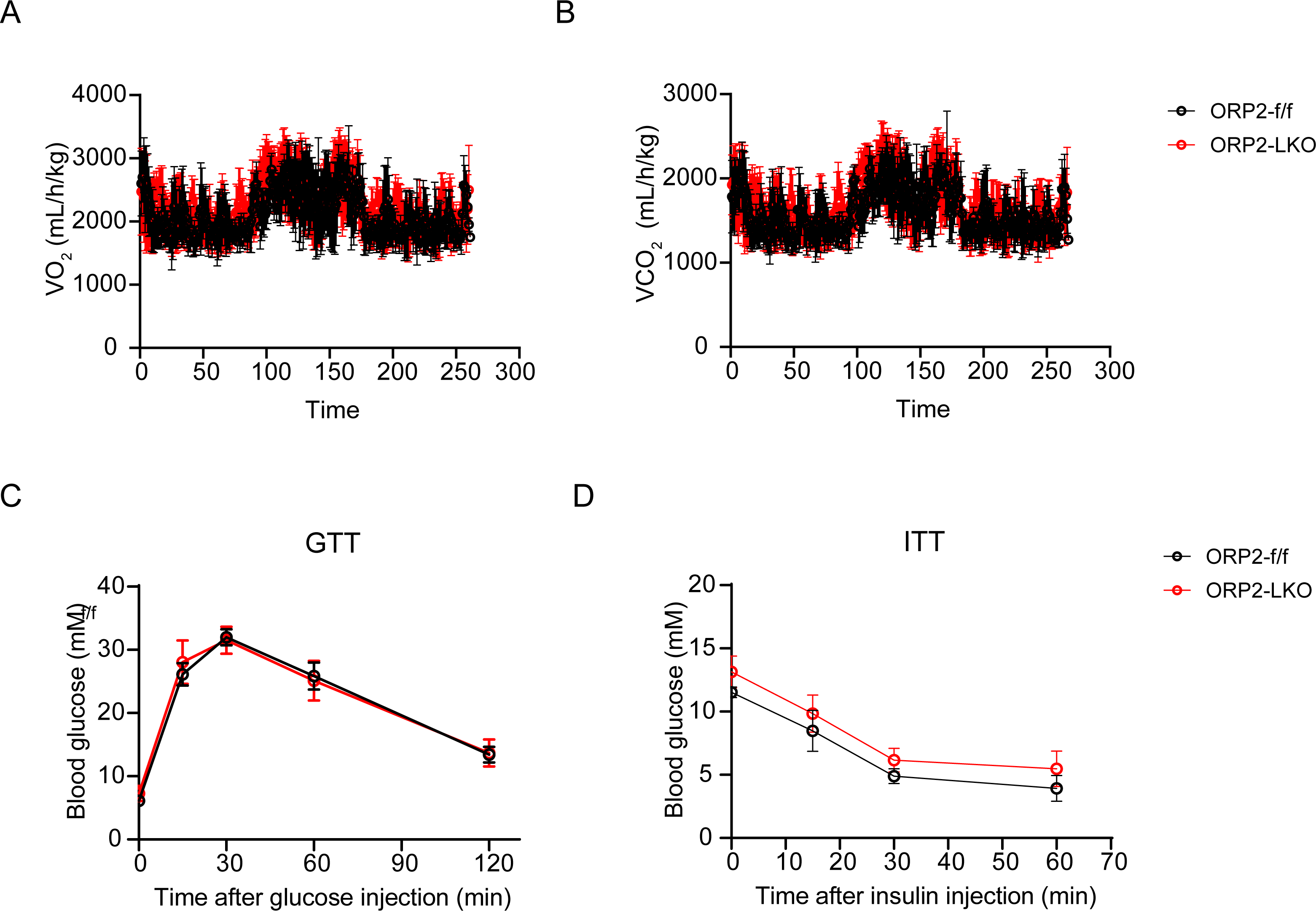
Systemic metabolic homeostasis, glucose tolerance and insulin sensitivity of ORP2-LKO mice on high fat diet. A and B. VO2 and VCO2 of mice subjected to metabolic cage analysis (n=5-9). C and D. Glucose tolerance test and insulin tolerance test of ORP2^f/f^ mice and ORP2-LKO mice (n=8-12). Data are presented as mean ± SD. Statistical significance was determined using a two-tailed Student’s t-test. *P < 0.05, **P < 0.01, ***P < 0.001.

